# Stress vulnerability promotes an alcohol prone phenotype in a preclinical model of sustained depression

**DOI:** 10.1101/358606

**Authors:** Danai Riga, Leanne JM Schmitz, Yvar van Mourik, Witte JG Hoogendijk, Taco J De Vries, August B Smit, Sabine Spijker

**Affiliations:** Department of Molecular and Cellular Neurobiology, Center for Neurogenomics and Cognitive Research, Amsterdam Neuroscience, VU University, Amsterdam, The Netherlands; Department of Anatomy and Neurosciences, Amsterdam Neuroscience, VU University Medical Center, Amsterdam, The Netherlands; Department of Psychiatry, Erasmus Medical Center, Rotterdam, The Netherlands

**Keywords:** depression susceptibility, depression resilience, comorbidity

## Abstract

Major depression and alcohol-related disorders frequently co-occur. Depression severity weighs on the magnitude and persistence of comorbid alcohol use disorder (AUD), with severe implications for disease prognosis. Here, we investigated whether depression vulnerability drives propensity to AUD at the preclinical level. We used the social defeat-induced persistent stress (SDPS) model of chronic depression in combination with operant alcohol self-administration (SA). Male Wistar rats were subjected to social defeat (5 episodes) and prolonged social isolation (~12 weeks) and subsequently classified as SDPS-prone or SDPS-resilient based on their affective and cognitive performance. Using an operant alcohol SA paradigm, acquisition, motivation, extinction and cue-induced reinstatement of alcohol-seeking were examined in the two subpopulations. SDPS-prone animals showed increased alcohol SA, excessive motivation to acquire alcohol, persistent alcohol-seeking despite alcohol unavailability, extinction resistance and increased cue-induced relapse; the latter could be blocked by the α2 adrenoreceptor agonist guanfacine. In SDPS-resilient rats, prior exposure to social defeat increased alcohol SA without affecting any other measures of alcohol-seeking and -taking. Our data revealed that depression proneness confers vulnerability to alcohol, emulating patterns of alcohol dependence seen in human addicts, and that depression resilience to a large extent protects from the development of AUD-like phenotypes. Furthermore, our data suggest that stress exposure alone, independently of depressive symptoms, alters alcohol intake in the long-term.

## Introduction

Major Depressive Disorder (MDD) is characterized by i) persistent negative mood, ii) loss of interest or inability to experience pleasure (anhedonia) and iii) mild cognitive impairment^1^. MDD is amongst the most detrimental psychiatric disorders, due to its high prevalence, substantial health burden and limited treatment response^2,3^. MDD commonly co-occurs with alcohol use disorder (AUD)^4^, defined by extreme alcohol preoccupation, alcohol craving and, recurrent episodes of relapse to alcohol use^1^, complicating its clinical profile and treatment^5, 6^. Approximately 1 out of 5 individuals diagnosed with MDD also suffers from AUD, a 4-fold incidence increase *vs.* healthy individuals^4^. In the majority of comorbid cases, MDD precedes the onset of alcohol dependence^7^. Notably, in epidemiological studies, the duration and severity of primary MDD appears to be a risk factor for developing secondary AUD^7^. Furthermore, comorbidity with MDD predicts greater severity of alcohol dependence^7^.

Exposure to severe and/or repeated stress is a well-established trigger of depressive symptoms, as observed both at the clinical^8, 9^ and the preclinical^10,11^ level. Response to stress determines the extent of depressive symptoms, and this is substantiated by an accumulating body of preclinical data examining individual variability to the effects of stress^12–14^. Notably, susceptibility to stress is characterized by dysregulation of the brain reward pathways^15–17^ and is accompanied by severe reward-associated behavioral deficits^18^, ^19^. For example, stress-susceptible animals display facilitation of drug-seeking behaviors, as observed in increased alcohol, amphetamine and cocaine intake^20–22^ and sensitization to the effects of cocaine and amphetamine^18^, ^23^.

Together, clinical and preclinical data support the interplay between the individual response to stress, depression severity and subsequent vulnerability to substance use disorder. Previously, we developed a rat paradigm that models primary depression and secondary AUD. Using social defeat-induced persistent stress (SDPS), we demonstrated that animals displaying a sustained depressive-like state showed enhanced vulnerability to alcohol-taking and -seeking, as reflected in excessive motivation to consume alcohol and heightened relapse rate^24^.

In the present study, we investigated whether individual variability to the effects of SDPS is associated with subsequent vulnerability to alcohol, and whether resilience to the effects of SDPS protects from the development of an addiction prone phenotype in the months following this stressor. In particular, we measured i) alcohol preference and consumption, ii) motivation for alcohol-taking, iii) persistence of alcohol-seeking during periods of unavailability, iv) extinction resistance and finally, v) reinstatement of alcohol-seeking behaviors in animals prone to the effects of SPDS and their resilient counterparts.

## Materials and Methods

### Animals & social defeat-induced persistent stress (SDPS)

Pair-housed male Wistar rats (Harlan CPB, Horst, Netherlands) 6-7 weeks old, weighing <200 g upon arrival were habituated (2 weeks), and exposed to social defeat-induced persistent stress (SDPS) followed by an operant alcohol self-administration (SA) paradigm, as previously described^24, 25^. In brief, SDPS animals (n=48) were subjected to five 15-minute daily social defeat sessions, based on the resident-intruder protocol. Rats were transported to the resident housing room and placed inside a resident cage (defeat cage). A transparent, perforated Plexiglas partition wall was used to separate the residents from the intruders, allowing for sensory exchange, but not for physical contact (pre-fight phase, 5 minutes). The wall was removed and Wistar rats were then exposed to a 5-minute fight phase, during which they were forced into submission. The defeat session concluded with an additional 5-minute period, during which the partition wall was placed back, separating the resident from the intruder (post-fight phase). A different resident was matched to each Wistar rat per day. Control rats (n=32) were exposed to an empty defeat cage, once per day for a total of 5 days. From the first defeat session or empty cage exposure onwards, all animals were single-housed and remained in social isolation for the rest of the experimental conditions, in absence of further sensory interaction with the stressor (residents). All experimental manipulations were conducted during the dark phase of a reversed 12-h light-dark cycle (lights on at 19.00 h). For the whole experimental period, animals received food and water *ad libitum.* All experiments were approved by the VU University Amsterdam Animal Users Care Committee.

### Selection of SDPS-prone *vs.* -resilient groups

SDPS rats were assigned to either SDPS-prone or -resilient subgroups following a two-step cluster analysis of affective and cognitive performance using the Schwarz’s Bayesian criterion^19^. In particular, rats were clustered based on their individual scores in the social approach-avoidance (SAA) and in object place recognition (OPR) tasks, assessed in weeks 5 and 9 post-defeat (for details see Supplemental Information, Fig. S1 and ^26^). From the emerging SDPS-prone and -resilient groups, data obtained from 10 rats (n=5 per subgroup) were used to describe alcohol-related effects of SDPS in the general population^24^, thus were not included in the alcohol SA analysis presented here. Control animals were divided in two equally performing groups (balanced average performance in SAA and OPR tests) and a total of 16 control rats participated in the experiments described below.

### Alcohol exposure

*Home cage consumption* - All animals were habituated to alcohol consumption using the two-bottle free / limited-access paradigm as previously described^27^. In brief, rats were exposed to gradually elevating alcohol concentrations (2-12% v/v) in the home cage for a total of 5 weeks. During the first 3 weeks of habituation in the home cage, alcohol was allowed for 24-h, followed by an alcohol free day before the next concentration increment. During the last 2 weeks, alcohol availability was limited to 1-h/day, to prime rats to the subsequent 1-h self-administration sessions. Water bottles were presented in parallel with alcohol, and were used to estimate alcohol preference *vs.* total liquid consumption. The position of alcohol and water bottles was alternated between days / sessions to avoid development of preference.

*Alcohol SA-Fixed Ratio* - Rats were trained to nose-poke for a 0.20 mL 12% alcohol reward in 1-h sessions given every other day. Alcohol delivery (US) was accompanied by discrete audiovisual stimuli (CS, 4-s active hole illumination and tone presentation) and was followed by a 15-s time-out period, during which nose-poking has no programmed consequences (alcohol unavailability period). Different reinforcement schedules (fixed ratio, FR) were used (FR1-3). In total, animals were subjected to 15 FR1, 5 FR2 and 5 FR3 sessions. Each FR increment was implemented after animals had reached stable performance, i.e., when there were no significant differences in responding between the last two sessions of each reinforcement schedule.

*Alcohol SA-Progressive Ratio* - Animals were subjected to five 2-h progressive ratio (PR) sessions, during which the effort (number of nose-pokes) to obtain a reward was progressively increased according to: response ratio = (5e^(02^ * ^reward number^))_5, rounded to the nearest integer.

*Alcohol SA - Time-Out performance -* Following PR, rats were re-trained to FR1 schedule (13 1-h sessions), to minimize between-group differences that could affect subsequent analysis of extinction performance. To decipher SDPS effects on alcohol-seeking during periods of unavailability, 4 of these FR1 sessions (sessions 4-7) included a doubled time-out interval (30-s).

*Alcohol SA-Extinction and Relapse* - Extinction training consisted of 1-h exposure to the training context in absence of alcohol and alcohol-associated cues. Following 15 daily sessions, operant responding was successfully extinguished (<6 active responses session) and all animals participated in two 30-minute cue-induced reinstatement sessions, at the start of which a single 0.20 mL alcohol reward was delivered. The two relapse tests were given with a 72-h interval and no additional extinction training, using a cross-over design (for details see Supplemental Information). This design was implemented to examine i) whether individual variability to the effects of SDPS alters reinstatement of alcohol-seeking and ii) whether guanfacine could prevent heightened relapse after SDPS, as shown before^24^.

### Statistical analyses

All behavioral data during alcohol SA, including FR, PR, extinction and relapse, were analyzed using repeated measures analysis of variance (ANOVA). When *P*-values reached level of significance (*P*<0.05), further analysis was performed using one-way ANOVA, paired or unpaired student’s t-test and post-hoc Tukey-HSD multiple comparisons. Homogeneity of variance was estimated and Hyunh-Feldt correction or non-parametric Kruskal-Wallis H test were implemented in case of assumption violation. All statistics were performed using IBM SPSS Statistics 24. In the alcohol SA paradigm, one animal (control) was excluded from statistical analysis as behavioral outlier (>2×SD from mean) in >50% of the FR3 and >50% of the PR sessions.

## Results

### Effects of SDPS on affective state and cognition

<u>Selection of SDPS-prone and -resilient groups</u> - Following two-step cluster analysis, two divergent groups were identified, as reflected by their performance in repeated SAA and OPR tests over a period of 9 weeks after exposure to social defeat (Fig. S1 and ^26^). The SDPS-resilient population coped with defeat and isolation stress and did not develop any of the affective or cognitive deficits commonly seen after SDPS^24, 25, 28^ In contrast, the SDPS-prone population showed long-lasting deterioration of affective performance, reflected in social withdrawal, accompanied by severe impairments in spatial memory. At week 12 after defeat, animals proceeded to the alcohol paradigm (Fig. 1a).

**Figure 1.**
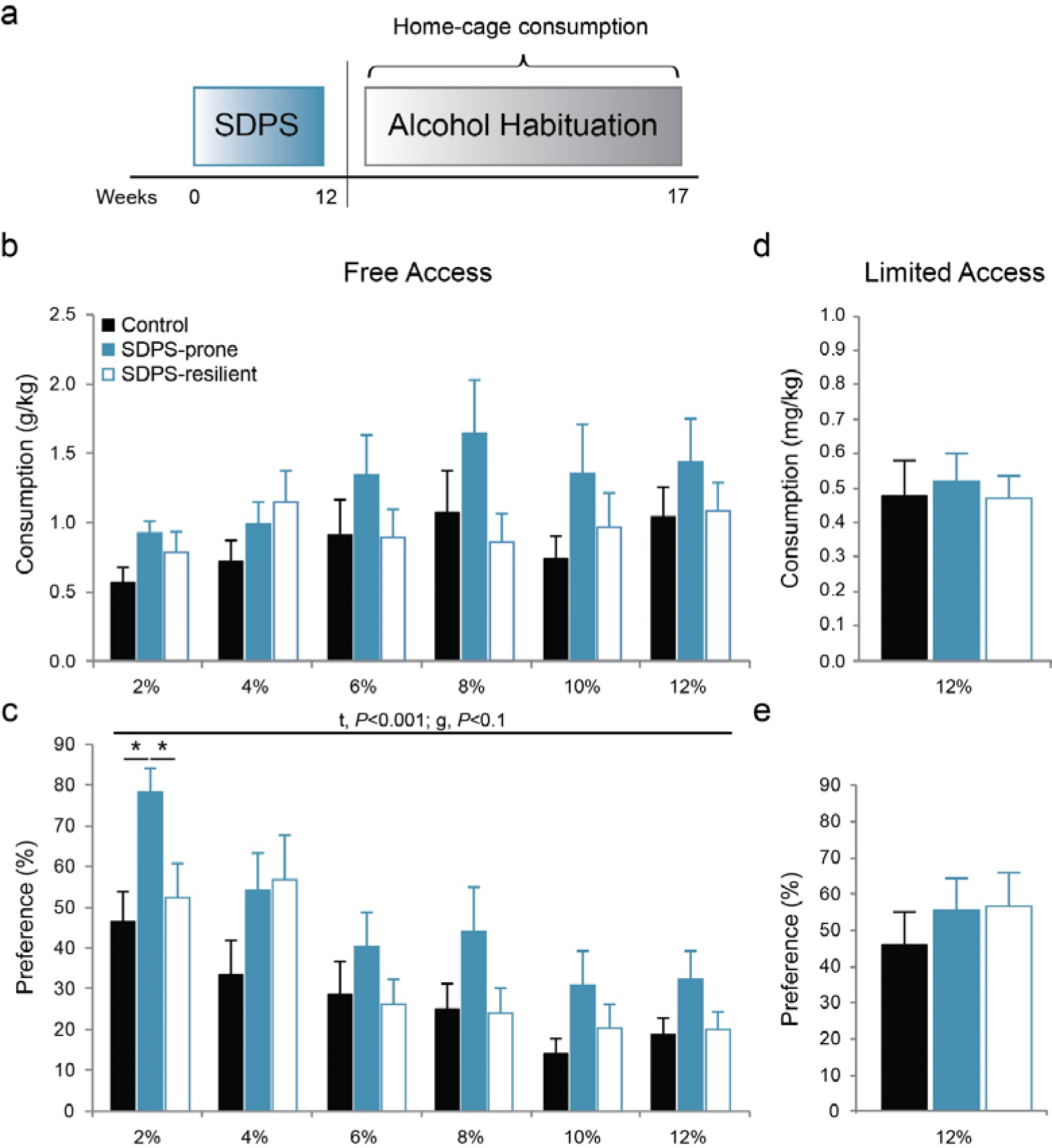
SDPS vulnerability increases preference for alcohol. **a)** Control rats and the two SDPS groups were habituated to progressively increasing concentrations of alcohol (2-12%) in the home cage, for a period of 5 weeks, using a two-bottle paradigm. **b)** Consumption of alcohol during 24-h free access, normalized for weight, revealed that, starting from 6% onwards, SDPS-prone rats displayed a relative increase in consumption of alcohol, albeit that no significant overall differences were observed between the 3 groups. **c)** Analysis of alcohol preference over water during 24-h free access, depicted as percentage of alcohol/total liquid consumed. A clear facilitation of alcohol consumption selectively in the SDPS-prone rats was observed. Between-group differences were most prominent at 2% alcohol concentration. SDPS-resilient and control groups showed similar preference for alcohol in all concentrations provided. **d)** Consumption of 12% alcohol during1-h limited access (average of 11 sessions). No difference in alcohol intake was observed between the 3 groups. **e)** Similarly, no between-group difference in preference for the alcohol solution was observed, as all 3 groups drank similar amounts of alcohol vs. water during the 1-h sessions. Repeated measures ANOVA main time (t) and group (g) effects are indicated; one-way ANOVA post-hoc group comparisons are depicted (c); \**P*<0.05.

### Effects of SDPS on alcohol-taking and -seeking

<u>Acquisition of operant alcohol self-administration</u> - During the 24-h free-access schedule in the home cage (Fig. 1a), similar alcohol consumption between control, SDPS-prone and SDPS-resilient animals was observed (Fig. 1b). By the end of the free-access period, all three groups consumed ~1 g/kg of 12% alcohol (control, 1.04±0.2; SDPS-prone, 1.44±0.3; and SDPS-resilient, 1.09±0.2 g/kg). Analysis of preference for the alcohol over the water solution during the entire free-access period showed a significant effect of alcohol (repeated measures ANOVA, F_EtoH_(3.55,113.62)=25.94, P<0.001, Fig. 1c), with no alcohol × group interaction (F_Etoh_ × GROUPH (7.10,113.62)=1.55, P=0.155) and a trend for between-group effects (F_GROUPH_ (2,32)=2.69, *P*=0.083). One-way ANOVA per alcohol concentration revealed that SDPS-prone rats preferred the 2% alcohol solution (F2%(2,34)=4.92, *P*=0.014) when compared to both control (*P*=0.005) and SDPS-resilient (*P*=0.030) groups. The SDPS-prone group showed a modest preference for alcohol in all concentrations examined, however no other statistical significant differences were observed.

During the subsequent 1-h limited access schedule no significant between-group effect was observed in either absolute consumption (F(2,34)=0.07, P=0.933, Fig. 1d) or preference (F(2,34)=0.52, P=0.597) for the 12% alcohol solution (average of 10 days, Fig. 1e). Together, SDPS-prone animals showed a moderate propensity towards passive alcohol intake that developed at >12 weeks from the last defeat exposure.

Following home cage alcohol habituation, animals were subjected to operant alcohol self-administration (Fig. 2a). Already in the first SA session, animals learned to discriminate between the active and the inactive hole, preferring the alcohol-associated one: Paired t-test, FR1active *vs.* FR1inactive controls, t(14)=5.95, *P*<0.001; SDPS-prone, t(9)=3.45, *P*=0.007; and SDPS-resilient, t(9)=2.77, *P*=0.022 (Fig. 2b, Fig. S2). Analysis of active responding during the three FR (1-3) reinforcement schedules, revealed an overall effect of training, representing an increase in responding following each change in schedule. Next, an overall effect of SDPS was observed for all 3 training ratios, and socially defeated animals, independently of subgroup, displayed increased number of responses in comparison with controls, as observed previously^24^. No training × group interaction effect was seen in any of the FRs tested, and no differences between SDPS-prone and SDPS-resilient animals were observed (for effects per FR schedule and post-hoc group comparisons see Table S1).

**Figure 2.**
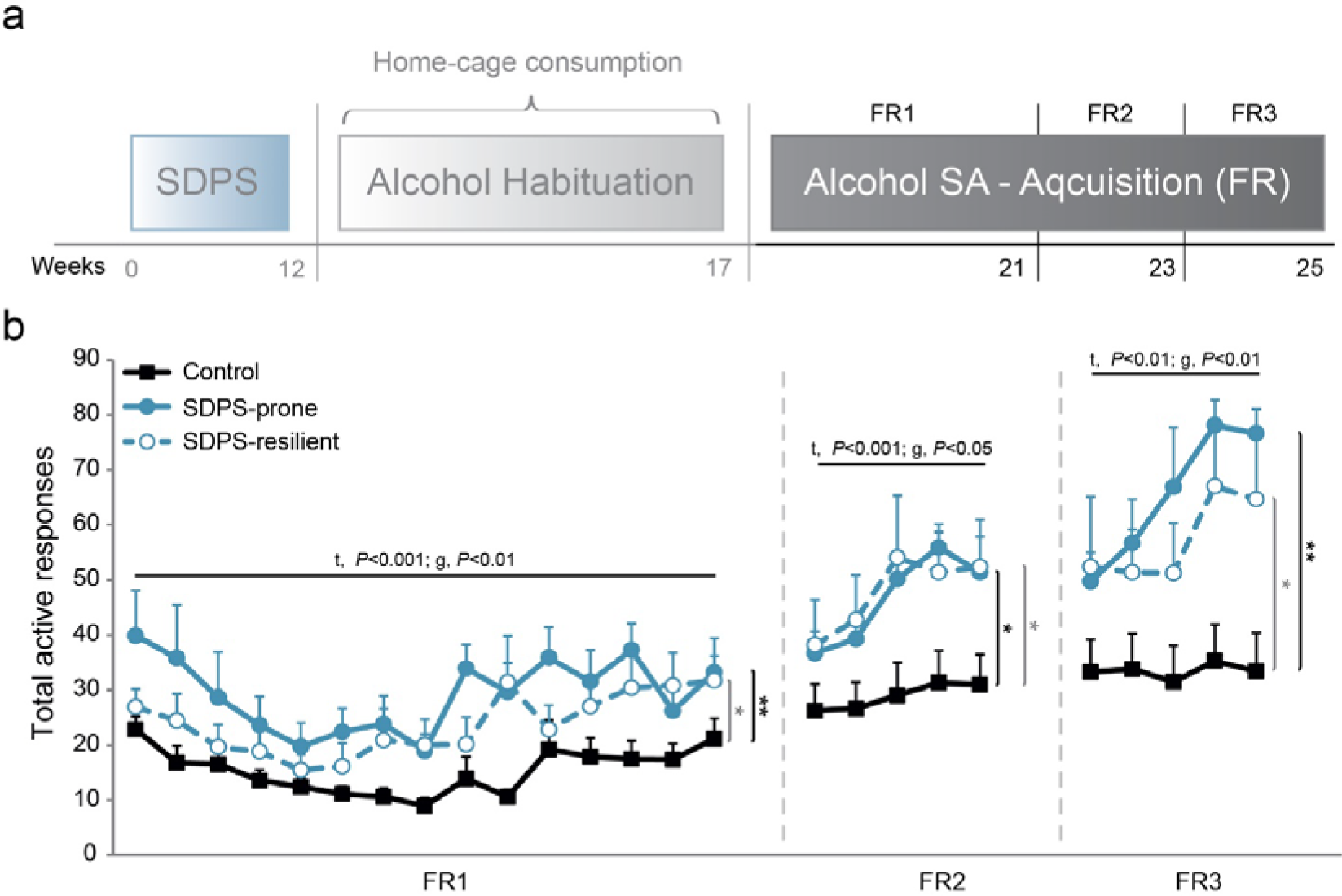
SDPS facilitates acquisition of operant alcohol self-administration. **a)** At ~4 months from the last defeat episode and following alcohol habituation at the home cage, all rats were subjected to a cue-coupled alcohol self-administration paradigm, starting with acquisition at FR. Different reinforcement schedules were used (FR1-FR3). **b)** Analysis of the number of responses to the active, alcohol-delivering hole during FR1 revealed significant training and group effects, as both SDPS groups displayed increased responding as compared with controls. Similarly, during FR2 and FR3 training schedules, the two SDPS groups exhibited enhanced responding for alcohol vs. controls. Although SDPS-prone animals showed relatively higher response rates, no group difference between the two SDPS groups was observed. Repeated measures ANOVA across the 3 reinforcement schedules, main time (t) and group (g) effects are depicted; pairwise group comparisons are indicated (vertical lines, black, SDPS-prone *vs.* controls; grey, SDPS-resilient *vs.* controls); \**P*<0.05; \*\**P*<0.01.

Analysis of the alcohol consumption data led to similar results, as in all 3 reinforcement schedules, the two SDPS groups gained higher number of rewards in comparison with controls (Fig. S2, Table S2). The average number of inactive responses per session was similar between the three groups in all reinforcement schedules given, supporting the view that task responding was alcohol-specific and excluding general psychomotor deficits long-term following social defeat (Fig. S2, Table S3). Together, the FR1-3 acquisition data reflected an SDPS-driven escalation of responding for an alcohol reward, which persisted, and was even exaggerated under more demanding reinforcement schedules.

<u>Progressive ratio (PR)</u> - After the last FR session we implemented PR training to study whether a similar increase in demand of reinforcement was evident after SDPS as observed previously^24^· ^25^ (Fig. 3a). Analysis over the 5 PR sessions showed no effect of training for the number of active responses (repeated measures ANOVA: F_PR_ (3.40,108.84)=2.07, *P*=0.100, Fig. 3b). A significant group effect was observed (F_GROUPH_ (2,32)=3.42, *P*=0.045), in absence of training × group interaction (F_PR×GROUPH_ (6.80,108.84)=0.72, *P*=0.653). Pairwise comparisons revealed that the SDPS-prone group showed a significantly higher number of responses for the alcohol reward *vs.* controls (*P*=0.016). No overall difference in responding between SDPS-resilient and control animals (*P*=0.128), nor between the two SDPS groups (*P*=0.375), was detected. Accordingly, a significant group effect in break points over the 5 PR sessions was observed (repeated measures ANOVA F_GROUPH_ (2,32)=3.44, *P*=0.044, Fig. 3c). Post-hoc comparisons revealed that this effect was driven by a strong increase in break points displayed by the SDPS-prone animals (*P*=0.016 *vs.* control), an effect that was absent in SDPS-resilient rats (*P*=0.121 *vs.* control). No difference in break points between the two SDPS groups was observed (*P*=0.388). Taken together, PR data confirmed that SDPS enhances motivation for alcohol-seeking^24^ and suggested that, to a large extent, SDPS-resilience prevents these motivational deficits.

**Figure 3.**
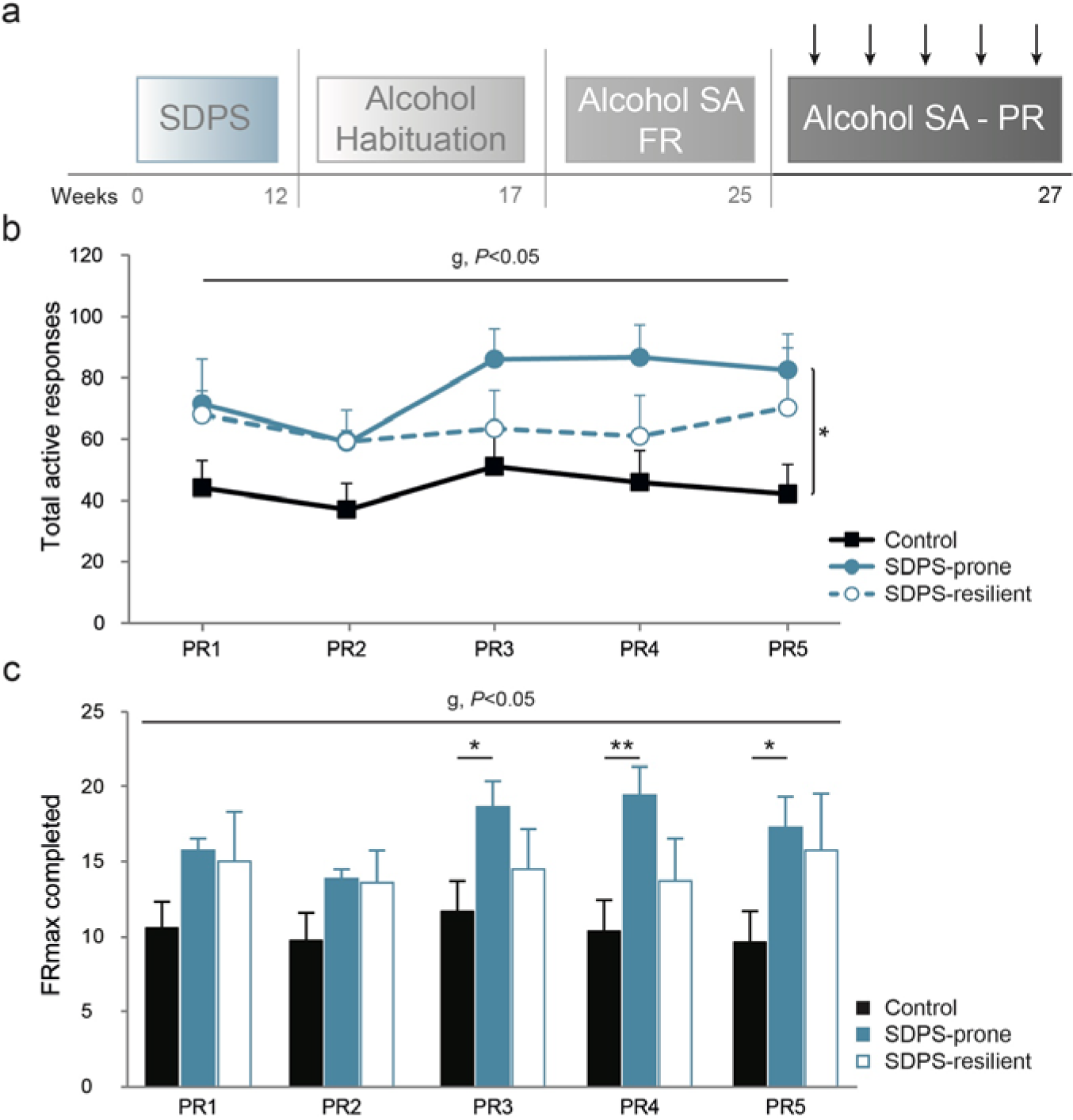
SDPS vulnerability increases motivation for alcohol intake. **a)** Following acquisition of alcohol SA, all animals were subjected to 5 progressive ratio sessions in which motivation for alcohol was assessed. **b)** Analysis of active responding revealed a main group effect, as SDPS-prone animals displayed significantly higher number of responses vs. controls. No difference between the two SDPS groups, or between the SDPS-resilient and control animals was observed. **c)** Similarly, break points (maximum FR reached) confirmed an SDPS-induced increase in motivation for alcohol. Importantly, this effect was seen only in the SDPS-prone rats, as SDPS-resilient animals did not differ from controls. Repeated measures ANOVA across the 5 PR sessions main group (g) effect and pairwise group comparisons are depicted (vertical lines, b); one way ANOVA main group (g) effects are indicated (c); \**P*<0.05; \*\**P*<0.01.

<u>Re-training on FR1</u> - Following PR, all animals were subjected to FR1 re-training (13 1-h sessions; reFR1) in order to normalize pre-existing group differences at the start of extinction (Fig. 4a). Repeated measures ANOVA revealed a significant training effect (: F_reFR1_ (4.54,145.43)=9.18, *P*<0.001) and no group × training interaction (F_reFR1xGROUp_(9.09,145.43)=1.14, *P*=0.335) as all animals gradually reduced their responding for an alcohol reward (Fig. S3). A significant group effect (F_GROUP_ (2,32)=5.83, *P*=0.007) pointed towards differential group performance over time. Post-hoc analysis further confirmed that, similar to acquisition in FRI, SDPS-prone rats showed enhanced responses compared with controls (*P*=0.002). This effect was not seen in SDPS-resilient animals (*P*=0.124 *vs.* controls). No differences between SDPS-prone and -resilient groups were detected (*P*=0.105). Notably, an initial carry-over effect in responding after PR was observed in the SDPS-prone group, which displayed higher number of active responses when compared with both control (*P*=0.001) and SDPS-resilient (*P*=0.018) groups at the first reFR1 session. Together, re-training in FR1 further indicated a stable long-term (~5 months after defeat) SDPS-triggered increase in alcohol-taking that was more prominent in the SDPS-prone rats.

**Figure 4.**
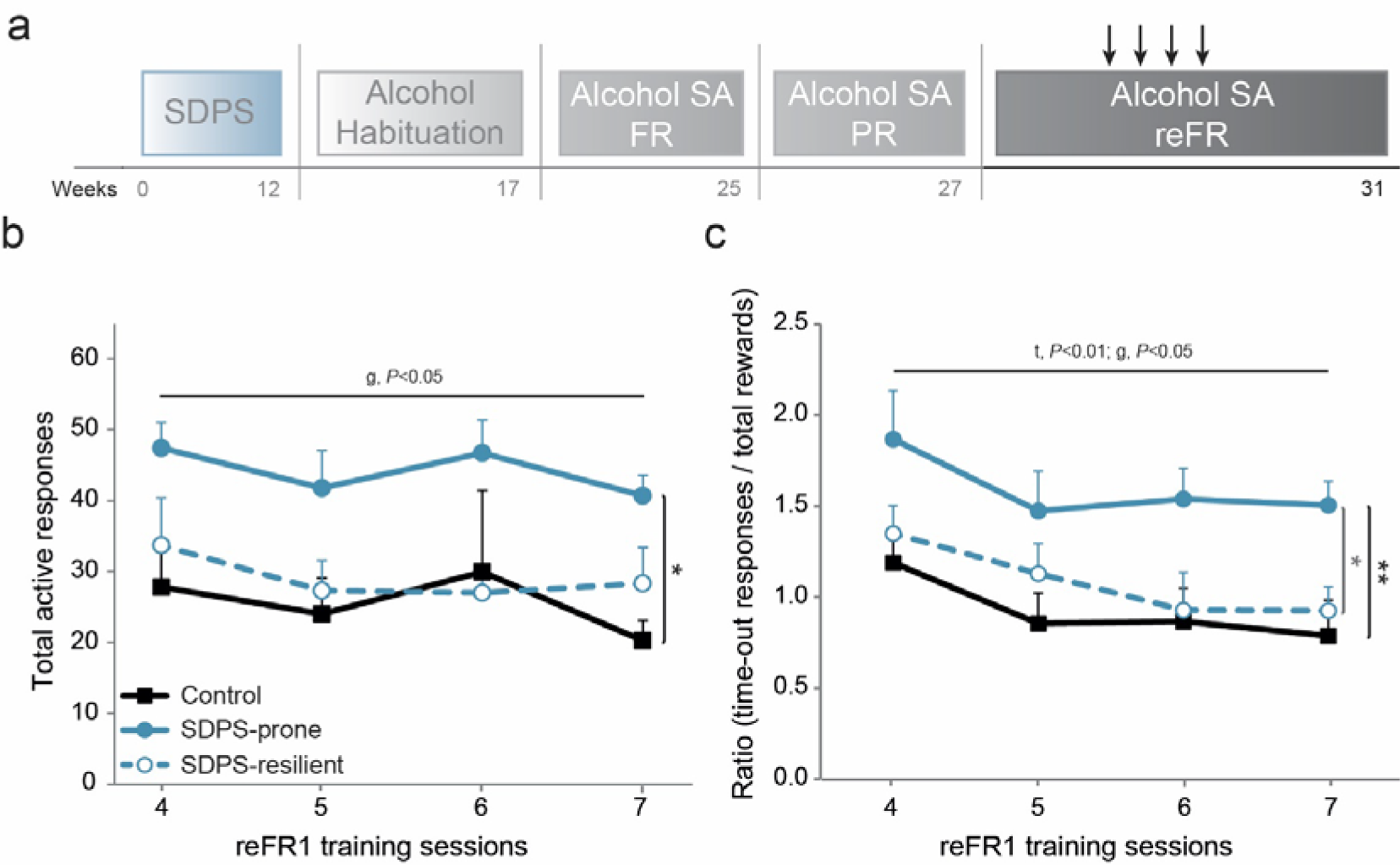
SDPS vulnerability induces persistent alcohol-seeking. **a)** After PR training, all animals were subjected to re-training in FR1 for 13 1-h sessions *(cf.* Fig. S3). During sessions 4-7, a 30-s time-out interval was implemented following each alcohol reward, doubling the original time-out period. **b)** Analysis of the active responses over these 4 sessions revealed a main group effect, as SDPS-prone rats displayed significantly increased responses *vs.* controls, and a trend *vs.* SDPS-resilient rats. No difference between SDPS-resilient and control groups was observed. **c)** To correct for pre-existing group differences in the chance of time-out responding, the ratio of time-out responses to actual rewards was calculated. Analysis of the time-out/reward ratio over 4 reFR1 sessions showed that the SDPS-prone group reached significantly higher ratio, when compared to both control and SDPS-resilient groups. No difference between control and SDPS-resilient groups was observed. Repeated measures ANOVA across the 4 reFR1 sessions main time (t) and group (g) effects and pairwise group comparisons (vertical lines, black, SDPS-prone vs. controls; grey, vs. SDPS-resilient) are indicated; \**P*<0.05; \*\**P*<0.01.

<u>Time-out performance</u> - Initially during acquisition of alcohol SA (FR1-3), we observed an SDPS-induced increase in responding during time-out periods, in which reward delivery was omitted (Fig. S2, Table S4). This effect was predominantly observed in the SDPS-prone and, to a lesser extent, in the SDPS-resilient groups. To further dissect alcohol-seeking behavior during alcohol unavailability periods, we introduced a 30-s time-out interval following each reward, in re-FR1 sessions 4-7, Fig. 4a. Analysis of active responses showed no training effect (repeated measures ANOVA: F_reFR1_ (1.77,56.73)=1.56, *P*=0.220) and no training × group interaction (F_reFR1xGROUP_ (3.55,56.73)=0.35, *P*=0.824), suggesting that overall responding for alcohol was not affected by the change in the duration of the time-out period (Fig. 4b, Fig. S3). Notably, a significant group effect was observed (F_GROUP_ (2,32)=3.65, *P*=0.037) due to increased responding in the SDPS-prone group when compared with controls (*P*=0.013), and a trend vs. SDPS-resilient (*P*=0.061) rats. No group difference was detected between SDPS-resilient animals and controls (*P*=0.617).

SDPS-prone animals displayed exaggerated active responding, and thus gained higher number of rewards. Since each reward delivery was followed by an alcohol unavailability period, SDPS-prone rats were presented with higher chances to respond during time-out. To control for this pre-existing difference, we next examined the relationship between time-out responses and the number of actual rewards obtained. In particular, we analyzed the ratio between non-reinforced responses and total rewards gained under this 30-s time-out interval (Fig. 4c). This revealed a significant training (repeated measures ANOVA: F_RATIO_ (3,96)=6.08, P=0.001) and group (F_GROUP_ (2,32)=5.43, P=0.009) effect, in absence of a training × group interaction (F_RATIO×GROUP_ (6,96)=0.23, *P*=0.966). Pairwise comparisons showed that SDPS-prone rats exhibited an increased ratio of non-reinforced responses *vs.* total rewards compared with both control (*P*=0.003) and SDPS-resilient (*P*=0.030) animals. No difference between the two latter groups was seen (*P*=0.454). Together, prolongation of the time-out period, during which alcohol delivery is omitted, increased alcohol-seeking selectively in the SDPS-prone group.

<u>Extinction</u> - Extinction training took place following re-exposure to FR1 (Fig. 5a). First, analysis of overall extinction performance during the whole training period revealed a significant time effect in absence of time × group interaction (repeated measures ANOVA, FEXT(5.74,183.69)=21.66, *P*<0.001; FEXT×GROUP(11.48,183.69)=1.18, *P*=0.302), as active responding decreased in all groups (Fig. 5b). A significant main group effect was detected (repeated measures ANOVA, Fgroup(2,32)=7.21, P=0.003), driven by the increased responding of SDPS-prone rats when compared with controls (*P*=0.001) and their resilient counterparts (*P*=0.026). No difference in responding between the latter two groups was observed (*P*=0.229). To further dissect the temporal component of the observed variation in extinction performance between the three groups, active responding was analyzed in 3 bins of 5 extinction sessions, representing each week of training in-between no-training weekend days (Fig. 5b). Repeated measures ANOVA showed significant time and group effects for the first two extinction training bins, i.e., EXT1-5: SDPS-prone *vs.* control, *P*=0.001 and SDPS-resilient *vs.* control, *P*=0.229); EXT6-10: SDPS-prone *vs.* control, *P*=0.001; SDPS-resilient *vs.* control, *P*=0.097) (Table S5), confirming a delayed extinction learning displayed by the SDPS-prone rats. No between-group difference in the last training week (EXT11-15) was observed, indicating that by the end of extinction training period all three groups performed similarly, extinguishing their responding for an alcohol reward (Fig. 5b and Fig. S4). Together, extinction data indicated that SDPS led to an initial delay in extinction learning that was most pronounced in the SDPS-prone group. Overall, SDPS-proneness resulted in persistent responding despite alcohol unavailability, a behavioral aspect not seen in resilient animals.

**Figure 5.**
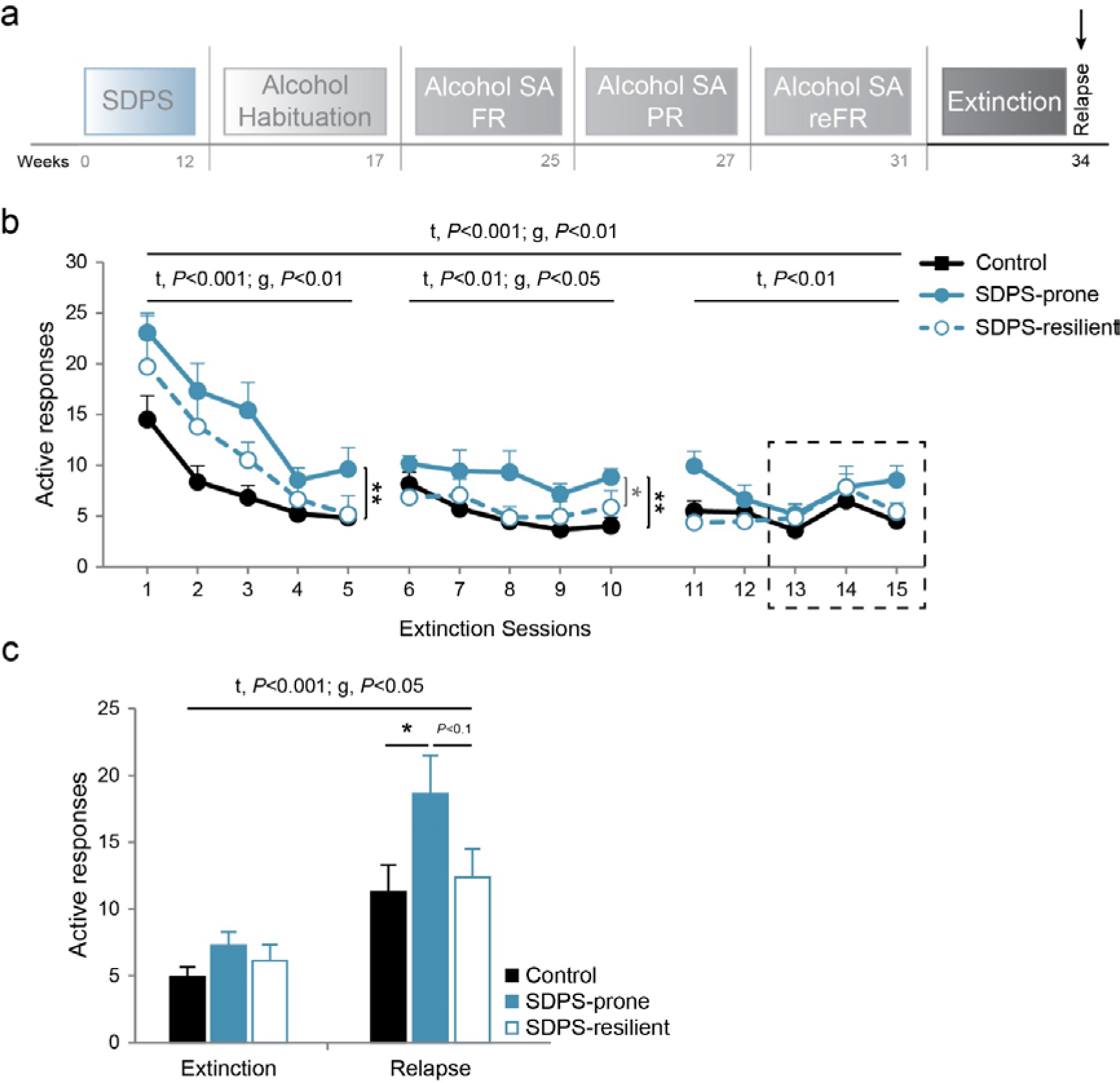
SDPS vulnerability delays extinction learning and facilitates reinstatement of alcohol-seeking. **a)** After re-training in FR1, all animals were provided with 15 1-h daily extinction sessions, during which uncoupling of the context (operant chambers) and the alcohol delivery was achieved. Following extinction training, all animals were subjected to cue-induced reinstatement, in absence of alcohol. **b)** Main training and group effects indicated differential extinction performance across the training period and among the three groups. Analysis of responding per training bin (3×5 extinction sessions) showed that the SDPS-prone group exhibited delayed extinction learning in the first two weeks (sessions 1-5 and 6-10), as reflected in increased active responses vs. controls and SDPS-resilient rats. Controls and SDPS-resilient animals responded similarly, illustrating that the observed effects on extinction were SDPS prone-specific. Analysis of the remaining extinction sessions (11-15) showed that all 3 groups were successfully extinguished by the end of the training period. **c)** Presentation of cues previously associated with reward delivery reinstated alcohol-seeking in all 3 groups, as compared to their average responding during the last 3 extinction sessions (indicated in b, dashed square). SDPS-prone animals showed increased relapse when compared with controls. No between-group differences in SDPS-resilient vs. control or SDPS-prone vs. SDPS-resilient groups were observed. Repeated measures ANOVA main time (t) and group (g) effects (b,c) and pairwise group comparisons (vertical lines, black, SDPS-prone vs. controls; grey, vs. SDPS-resilient, b) are indicated; One-way ANOVA main group effect is indicated (c); \**P*<0.05; \*\**P*<0.01.

<u>Cue-induced reinstatement</u> - Following extinction of the context of alcohol delivery, two cue-induced reinstatement tests were implemented in order to examine i) the effects of individual SDPS variability to alcohol relapse (Fig. 5a, saline test) and ii) the efficacy of guanfacine pre-treatment in preventing SDPS-induced heightened relapse (Fig. S5, guanfacine test). The average number of active responses during the last 3 extinction sessions (EXT13-15; one-way ANOVA, F_GROUP_(2,34)=1.61, *P*=0.215) was compared to responses gained during the two relapse tests^24^. During the saline test, a significant relapse effect was detected following presentation of alcohol-associated cues, in absence of relapse × group interaction (repeated measures ANOVA: F_RELAPSE_(1,32)=30.72, *P*<0.001; F_RELAPSE×GROUP_(2,32)=1.26, *P*=0.296), as all animals increased responding compared with their extinction performance (paired t-test: controls, t(14)=-3.15; *P*=0.007; SDPS-prone, t(9)=-3.59; *P*=0.006; and SDPS-resilient, t(9)=-2.70; *P*=0.024, Fig. 5c). Notably, a main group effect was observed (repeated measures ANOVA: F_GROUP_(1,32)=3.74, *P*=0.035), which was due to increased relapse in SDPS-prone animals *vs.* control (*P*=0.011) and a trend *vs.* SDPS-resilient (*P*=0.072) groups. The two latter groups performed almost identical (*P*=0.520). Analysis of relapse performance alone confirmed that SDPS vulnerability triggered increased reinstatement of alcohol seeking (one-way ANOVA SDPS-prone, post-hoc: *P*=0.031 *vs.* controls; *P*=0.092 *vs.* SDPS-resilient group; and controls *vs.* SDPS-resilient, *P*=0.730). In agreement with our previous observation^24^, pre-treatment with guanfacine blocked reinstatement of alcohol-seeking in all three groups, and prevented increased reinstatement in the SDPS-prone subpopulation (Fig. S5). Taken together, relapse data pointed to an SDPS-induced facilitation of reinstatement of alcohol-seeking, as shown previously^24^. Importantly, this effect was selectively seen in the SDPS-prone individuals, whereas SDPS-resilience seemed to protect from an increase in relapse.

## Discussion

In the present preclinical study, we examined the effects of depression vulnerability on alcohol-seeking and -taking behavior. This was to establish whether SDPS-proneness, which is associated with primary depressive-like symptoms, promotes secondary alcohol use disorder, two phenotypes that are often comorbid in humans^2^. We used SDPS in rats to model a depressive-like state that is sustained for at least 6 months following exposure to stress^24,26^, mimicking chronic depression in humans. Our approach allowed for measuring different features of alcohol-seeking and -taking in depression-prone *vs*. -resilient individuals, drawing parallels to the human disorder.

In the population of patients diagnosed with recurrent depression, comorbid alcohol abuse reaches a striking 40%^2^, indicating common genetic and/or environmental causes. In agreement with this, we previously showed that SDPS, when coupled with operant alcohol self-administration, promotes AUD-like behaviors, as reflected in excessive motivation for alcohol and increased relapse rate^24^. Here, we extend these findings by showing that animals selected for depression susceptibility exhibit greater vulnerability to alcohol in terms of persistent seeking despite alcohol unavailability and delayed extinction of alcohol-related learning. Furthermore, in our model, depression resilience subdued the emergence of addiction proneness, including enhanced motivation, resistance to extinction and aggravated alcohol-seeking during unavailability periods (time-out and reinstatement). Of note, stress exposure resulted in a lasting increase in instrumental alcohol-taking in SDPS-resilient individuals, which showed no depressive-like affective and cognitive deficits.

### Exposure to SDPS precipitates alcohol-taking vulnerability

In humans, depression-susceptibility primes the development of an alcohol-vulnerable phenotype^4^, which is characterized by core manifestations of AUD, including increased intake^1^. We showed that in the home cage, SDPS-prone rats displayed preference for a low concentration of alcohol compared with controls, an effect not observed in the general population of SDPS rats^24^. This moderate preference for alcohol was stable across the different alcohol concentrations used, when alcohol was provided *ab libitum*. This phenomenon, selectively seen in rats that display severe depressive-like symptoms, could indicate an attempt for self-medication^29^, as it has been long hypothesized based on the anxiolytic properties of alcohol.

Preclinical literature supports detrimental effects of stress on alcohol consumption and alcohol-seeking, although there is a complex interplay between biological factors governing stress responses and the methodological variations in stress application and alcohol exposure^30, 31^. We previously reported that exposure to the SDPS paradigm facilitated acquisition of alcohol self-administration in demanding, fixed schedules of reinforcement in the general population^24^. Here, we replicated this finding, in fact showing that SDPS increases alcohol intake independently of the presence of depressive-like symptoms, namely, affective and cognitive deficits. In particular, we report that SDPS-resilient animals, which exhibit no difference in the SAA and OPR tasks as compared with controls^26^, showed increased alcohol acquisition during fixed-ratio responding.

This SDPS-induced facilitation of operant alcohol intake has important clinical implications as it indicates that exposure to brief but severe social stress in combination with prolonged, subthreshold stressors, e.g., social isolation can render an individual vulnerable to alcohol intake, independently of its measurable depressiogenic effects. This is in agreement with clinical studies implicating stress coping styles, i.e., an individual’s response to perceived stress, in the development of alcohol dependence^32^, ^33^. It is noteworthy that, although no significant difference between the two SDPS groups was observed during acquisition of self-administration, SDPS-prone rats showed a considerable increase in responding for alcohol compared with their resilient counterparts. This surfaced in particular when more effort was required to obtain an alcohol reward, i.e., during FR3, acting as prelude to PR performance.

### SDPS-susceptibility is accompanied by excessive motivation towards alcohol

In depression, anhedonia, including disruptions in normal anticipatory response and in goal-directed behavior, is considered a core symptom^34^ that has been employed to assess depression-susceptibility^19^. At the preclinical level, progressive ratio (PR) responding has been extensively used to dissect the effects of depressive-like state in motivation towards natural^25^ and drug-related rewards^35^. In drug addiction, persistent preoccupation and heightened motivation to acquire the drug of abuse are central to disease diagnosis^1^. Consequently, PR schedules have been employed to assess the incentive value of drugs of abuse both in humans and in rodents^36, 37^ and are considered essential in prediction of addiction-proneness at the preclinical level^38^.

We previously showed that SDPS dramatically increased alcohol break points and that the SDPS-induced depressive state, as manifested in social avoidance after defeat, was predictive of a high motivational drive to seek alcohol^24^. Here, we demonstrated that SDPS-prone rats showed a similar, yet exaggerated response to PR training, confirming the crucial interaction between depression-susceptibility and alcohol-vulnerability. In support of this, resilience to SDPS limited alcohol-related motivational overdrive, as SDPS-resilient rats demonstrated PR performance similar to controls. This further indicates the conducive role of depression susceptibility on AUD-like manifestations.

### SDPS-susceptibility elicits extinction-resistance

The depressive-like state was accompanied by resistant extinction learning, which carried on for the first two weeks of extinction training. In particular, SDPS-prone rats showed delayed discontinuation of responding, as compared both with controls and their resilient counterparts. This delayed incorporation of contextual updates corresponds well to overgeneralization of conditioned stimuli, as it is hypothesized in depression^39^. In favor of this notion, mice exposed to repeated social defeat stress display delayed fear extinction and exhibit generalization of fear^40^. Alternatively, delayed extinction performance could reflect SDPS-induced deficits in cognitive flexibility, as observed in depressed patients^41^. Impaired reversal learning, especially when it requires the inhibition of behavioral patterns driven by affective information, has been observed in the clinic^42^. At the preclinical level, exposure to social defeat during adolescence is associated with deficits in reversal learning during adulthood^43^.

Importantly, these deficits depend on social context, as they were reversed following social housing but maintained in adults that, similar to our paradigm, remained in social isolation^43^.

Depression-resilience limited the emergence of delayed extinction. Particularly, during the first week of extinction training, SDPS-resilient animals performed midway of the two other groups, mimicking the effect of SDPS in the general population^24^. From then onwards, extinction responding in SDPS-resilient individuals mirrored the performance of control rats, indicating that in these animals, extinction of previously learned but currently inappropriate behavioral patterns is intact. This is in accordance with the notion that resilience to severe stress is characterized by facilitated extinction of non-relevant information^44^, i.e., adaptive extinction learning. Notably, facilitation of reversal learning, namely, a swift from learned responses towards the most adaptive ones, is observed following administration of tricyclic antidepressants^45^ and of selective serotonin reuptake inhibitors^46^. Thus it is possible that in SDPS-resilient animals, extinction learning is mediated via adaptations of the serotoninergic and noradrenergic systems that promote cognitive/ behavioral flexibility.

### SDPS-susceptibility prompts persistent alcohol-seeking and intensifies reinstatement

SDPS-prone animals exhibited escalated alcohol-seeking during time-out periods, after doubling the time interval before a subsequent alcohol reward was available. Our data point to an inability of SDPS-prone rats to withhold active responding, which led to premature behavioral responses, a hallmark of reduced inhibitory control. Increased premature^47^ and anticipatory^48^ responding are linked to alcohol abuse and dependence, while behavioral loss of control and cognitive impulsivity are associated with depression severity^49–51^. Our previous observations that guanfacine, a cognitive enhancer^52^ used against attention deficit-hyperactivity disorder (ADHD), ameliorates the effects of SDPS on alcohol^24^, further support dysregulated impulse control after SPDS. Together, it is possible that in the depression-prone population, behavioral disinhibition promoted compulsive-like alcohol-seeking. Future studies should address this possibility by subjecting SDPS-prone animals to putative measures of impulsive choice and action, such as the delayed reward discounting task. Likewise, it would be of interest to examine whether this phenotype is drug-specific or whether it is extended to non-drug natural rewards such as sucrose or food.

SDPS-proneness was accompanied by heightened reactivity to alcohol-signifying cues and promoted excessive reinstatement of alcohol-seeking, a common characteristic of animals displaying dependence-like behaviors^38^ and in SDPS-induced chronic depression^24, 25^ In accordance with the effect of SDPS susceptibility in timeout responding, aggravated relapse could reflect reduced cognitive control, i.e., an inability in refraining from alcohol-seeking and -taking. This notion is supported by our observations that guanfacine pretreatment normalized excessive alcohol-seeking in the SDPS-prone animals during reinstatement. Dysfunctional inhibitory control towards alcohol has been observed before in mice that, similar to the SDPS-prone group, show preference for alcohol^47^. Likewise, mice selected for decreased inhibitory control show escalated motivation for alcohol, increased time-out responding and enhanced relapse following presentation of alcohol signifying cues^53^, three phenotypes present in the SDPS-prone population. It is possible that these deficits are exaggerated by dysregulation of the brain’s reward system, which is present in the depression-susceptible but not in the unsusceptible subpopulation^18^, leading to maladaptive responsivity to alcohol-paired cues. Animals resilient to the effects of SDPS showed no changes in alcohol-seeking nor heightened relapse, further indicating that adaptability to adverse life events ameliorates the expression of addiction-like phenotypes^54^.

### Is SDPS-proneness a common denominator in depression- and addiction-vulnerability?

Currently, the cause and the temporal order of the development of comorbid MDD and AuD is lively debated^55^, highlighting the role of alcohol abuse as a risk factor for depression and vice versa^56^. In this discussion, genetic and environmental factors, as well as their interaction, are considered crucial for the emergence of comorbidity^57–59^. We report that individual variability to the effects of chronic stress, which precipitates or precludes the development of a chronic depressive-like state^26^, is related to the emergence or absence of AUD-like manifestations, respectively. In our model, maladaptive stress coping, which leads to propensity to primary depression, exaggerates secondary AUD-like behaviors. Notably, depression-resilience limits the emergence of the full comorbid phenotype, protecting from changes in extinction learning, persistent alcohol-seeking and relapse vulnerability. Our data might be explained by 1) a common (epi)genetic predisposition underlying the two diseases, 2) depression as a factor that confers vulnerability to alcohol abuse, 3) a combination of the two. For example, increased alcohol preference in the SDPS-prone individuals might reflect a genetic vulnerability to alcohol^60^ in the same individuals that show depression susceptibility^57^, especially when these individuals are exposed to adverse environmental conditions^61^, ^62^.

Collectively, our data support the hypothesis that depression susceptibility promotes addiction-like behaviors in the rat. It is worth mentioning that despite our systematic analysis of AUD-like phenotypes, we did not examine the effects of SDPS-proneness in resistance to punishment, which emulates continuation of drug use regardless of adverse consequences. Responding for alcohol, when its delivery is paired with an aversive stimulus, e.g., an electrical shock, is a robust measure of compulsive drug-seeking, used to identify alcohol-prone individuals^63^. Thus, it would be of great interest to evaluate the performance of the SDPS-prone population in this respect.

In conclusion, we showed that the SDPS model can be used to screen for depressed individuals with propensity to alcohol abuse and identify those that will develop AUD-like phenotypes. In turn, this can be used as a starting point for further research into (epi)genetic vulnerability factors, as well as molecular mechanisms that underlie the comorbid phenotype.

## Acknowledgements

ABS and SS received support from HEALTH-2009-2.1.2-1 EU-FP7 ‘SynSys’ (#242167); DR, LJMS and ABS received support from NBSIK PharmaPhenomics grant LSH framework FES0908; DR, LJMS and SS were supported by an NWO VICI grant (ALW-Vici 016.150.673 / 865.14.002).

## Authors Contribution

DR, SS: study concept & design

DR, LJMS, YvM: animal data acquisition

DR, LJMS: data analysis & interpretation

DR, SS: manuscript drafting

DR, SS, TJdV, WJGH, ABS: manuscript revision

